# Non-breeding waterbirds benefit from protected areas when adjusting their distribution to climate warming

**DOI:** 10.1101/2021.04.26.441480

**Authors:** Elie Gaget, Diego Pavón-Jordán, Alison Johnston, Aleksi Lehikoinen, Wesley M. Hochachka, Brett K. Sandercock, Alaaeldin Soultan, Hichem Azafzaf, Nadjiba Bendjedda, Taulant Bino, Luca Božič, Preben Clausen, Mohamed Dakki, Koen Devos, Cristi Domsa, Vitor Encarnação, Kiraz Erciyas-Yavuz, Sándor Faragó, Teresa Frost, Clemence Gaudard, Lívia Gosztonyi, Fredrik Haas, Menno Hornman, Tom Langendoen, Christina Ieronymidou, Vasiliy A. Kostyushin, Lesley J. Lewis, Svein-Håkon Lorentsen, Leho Luiujoe, Włodzimierz Meissner, Tibor Mikuska, Blas Molina, Zuzana Musilová, Viktor Natykanets, Jean-Yves Paquet, Nicky Petkov, Danae Portolou, Jozef Ridzoň, Samir Sayoud, Marko Šćiban, Laimonas Sniauksta, Antra Stīpniece, Nicolas Strebel, Norbert Teufelbauer, Goran Topić, Danka Uzunova, Andrej Vizi, Johannes Wahl, Marco Zenatello, Jon E. Brommer

## Abstract

Climate warming is driving changes in species distributions, although many species show a so-called climatic debt, where their range shifts lag behind the fast shift in temperature isoclines. Protected areas (PAs) may impact the rate of distribution changes both positively and negatively. At the cold edges of species distributions, PAs can facilitate species distribution changes by increasing the colonization required for distribution change. At the warm edges, PAs can mitigate the loss of species, by reducing the local extinction of vulnerable species. To assess the importance of PAs to affect species distribution change, we evaluated the changes in a non-breeding waterbird community as a response to temperature increase and PA status, using changes of species occurrence in the Western-Palearctic over 25 years (97 species, 7,071 sites, 39 countries, 1993– 2017). We used a community temperature index (CTI) framework based on species thermal affinities to investigate the species turn-over induced by temperature increase. In addition, we measured whether the thermal community adjustment was led by cold-dwelling species extinction and/or warm-dwelling species colonization, by modelling the change in standard deviation of the CTI (CTI_sd_). Using linear mixed-effects models, we investigated whether communities within PAs had lower climatic debt and different patterns of community change regarding the local PA surface. Thanks to the combined use of the CTI and CTI_sd_, we found that communities inside PAs had more species, higher colonization, lower extinction and the climatic debt was 16% lower than outside PAs. The results suggest the importance of PAs to facilitate warm-dwelling species colonization and attenuate cold-dwelling species extinction. The community adjustment was however not sufficiently fast to keep pace with the strong temperature increase in central and northeastern Western-Palearctic regions. Our study underlines the potential of the combined CTI and CTI_sd_ metrics to understand the colonization-extinction patterns driven by climate warming.

## Introduction

Global warming is one of the major causes of biological changes among the growing cocktail of anthropic pressures on the natural world (Monastersky 2014). There are several studies documenting global species distribution shifts towards the poles (Parmesan & Yohe 2003, Chen et al. 2011) which are driven by colonization at the leading distribution edge and extinction at the trailing edge (Thomas and Lennon 1999, Franco et al. 2006). However, distribution changes have mostly been insufficient to track the thermal isocline shifts, leading to climatic ‘debt’ in species distributions (Chen et al. 2011, Devictor et al. 2012). Furthermore, the pressures from climate change may be exacerbated by other factors interacting with colonization and extinction processes (Hill et al. 2001, Brook et al. 2008), like habitat fragmentation (Opdam and Wascher 2004, Hill et al. 2001) or land-use change (Auffret and Thomas 2019, Gaget et al. in press). However, some of these interactions may be positive, for example, protected areas may positively alter species ability to respond to climate change (Thomas et al. 2012).

Protected areas (hereafter, PAs) are expected to facilitate species distribution shifts in response to climate warming by reducing anthropic pressures on ecosystems (Monzón et al. 2011). PAs are one of the most efficient solutions to protect ecosystem of high biological importance (Godet and Devictor 2018). At the leading edge of species distributions, colonization may occur more likely in PAs (Hiley et al. 2013, Gillingham et al. 2015, Lehikoinen et al. 2019, Peach et al. 2019), particularly with large PA surface (Gaüzère et al. 2016), promoting range expansion (Thomas et al. 2012, Pavón-Jordán et al. 2015). Conversely, species extinction at the trailing edge can be reduced within PAs (Gillingham et al. 2015, Lehikoinen et al. 2019, Peach et al. 2019). In view of these contrasting patterns, it is important to evaluate in a comprehensive framework the effects of PAs on species distributions throughout the overall community of species.

Temperature driven shifts in species distributions will reshuffle community structure, with colonization of warm-dwelling species and/or extinction of cold-dwelling species (Devictor et al. 2008). Community adjustment to climate warming can be assessed with the intuitive community temperature index (hereafter, CTI), based on the average species thermal affinities in a community (Devictor et al. 2008). The CTI allows us to identify how local conditions such as site protection influence the community adjustment to warming (Gaüzère et al. 2016, Santangeli et al. 2017), and quantify any delay in tracking climate warming, namely the climatic debt (Devictor et al. 2012). In addition to the average community response measured with the CTI, the variance of the response provides a complementary indicator with which to investigate the species colonization-extinction processes (Fig. 1, Gaüzère et al. 2019).

**Figure 1:**
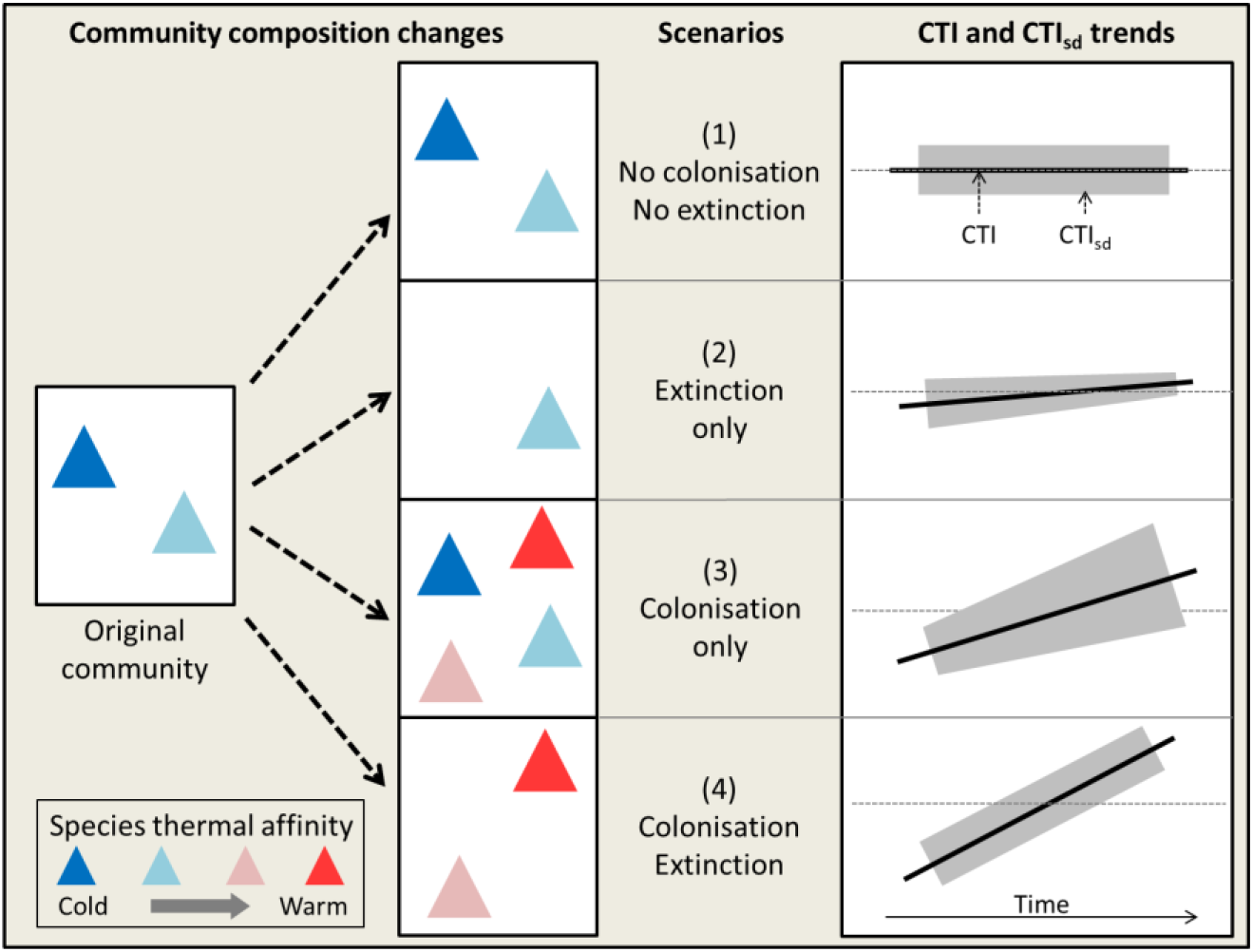
Schematic models of the four theoretical species colonization and/or extinction scenarios depending of their thermal affinities in response to climate warming and subsequent trends of community temperature index (CTI, i.e., thermal average) and CTI standard deviation (CTI_sd_, i.e., thermal standard deviation) over time (see Gauzere et al. 2019). Species are represented by colored triangles: blue to red correspond to cold- and warm-dwelling species, respectively. The different scenarios are, (1) ‘No colonization-No extinction’ causes no CTI and CTI_sd_ changes; (2) ‘Extinction only’ causes CTI increase and CTI_sd_ decrease by the loss of cold-dwelling species; (3) ‘Colonization only’ causes CTI and CTI_sd_ increase by the gain of warm-dwelling species; (4) ‘Colonization-Extinction’ causes CTI increase by the species thermal turn-over, but no CTI_sd_ directional change. The code for simulations is in Appendix 1.

Here, we investigated the community adjustment of non-breeding waterbirds to climate warming throughout the Western-Palearctic over 25 years and whether the patterns of change differed within and outside of PAs. This region, extending from the Mediterranean biodiversity hotspot to the Arctic, faces substantial anthropic pressures (IPCC 2014, IPBES 2018a, 2018b). Despite great conservation efforts, wetlands in this region have suffered drastic damages (Dixon et al. 2016) and many waterbird populations have been declining for decades (Gardner & Davidson 2011). Because of this, waterbirds have been targeted with a large-scale monitoring program, the International Waterbird Census (IWC, Delany 2010), which provides unique data to investigate the effectiveness of conservation strategies at continental scale (Pavón-Jordán et al. 2015, Amano et al. 2018). We expect that in response to climate warming, warm-dwelling waterbirds will colonize more in PAs and cold-dwelling species may be more resilient within PAs, as they usually contain good quality habitat (Lawson et al. 2014). Despite numerous studies on waterbird distribution changes in response to climate warming (e.g. Maclean et al. 2008, Lehikoinen et al. 2013, Pavón-Jordán et al. 2019), including conservation measures (Johnston et al. 2013, Pavón-Jordán et al. 2015, Gaget et al. 2018, Marion and Bergerot 2018), assessments of differences in waterbird distribution changes at community level inside and outside PAs are still lacking.

We analyzed an extensive dataset on waterbird occurrence (97 species) across 39 countries (7,071 sites), within the CTI framework (Devictor et al. 2008) and the related community thermal standard deviation (hereafter CTI_sd_, Fig. 1) to i) evaluate whether the community adjustment to climate warming was higher, and the climatic debt lower, inside PAs, ii) identify whether within PAs there are more colonization of warm-dwelling species and fewer extinction of cold-dwelling species, and iii) investigate whether the community adjustment to climate warming was improved where local PA surface was larger.

## Material and methods

### Study area and waterbird monitoring

We used International Waterbird Census (IWC) data from almost all of the Western-Palearctic (39 countries, Fig. 2, Appendix 2) from 1993–2017. The IWC monitors non-breeding waterbirds with a single count each year by ornithologists, professional or citizen scientists, in January and is coordinated by Wetlands International (www.wetlands.org, see Delany (2010) for the protocol). To ensure a long-term survey of community changes, we filtered the original data down to information from the 7,071 sites included in the study (Fig. 2) that each have at least five counts, with one count in each decade (1990s, 2000s and 2010s; 16.6 ± 5.6 counts per site) and at least two species per count (n = 117,325 counting events, Appendix 2). We included the 97 non-vagrant waterbird species overwintering in the Western-Palearctic (Appendix 3) listed in the African-Eurasian Migratory Waterbird Agreement (AEWA, http://www.unep-aewa.org).

**Figure 2:**
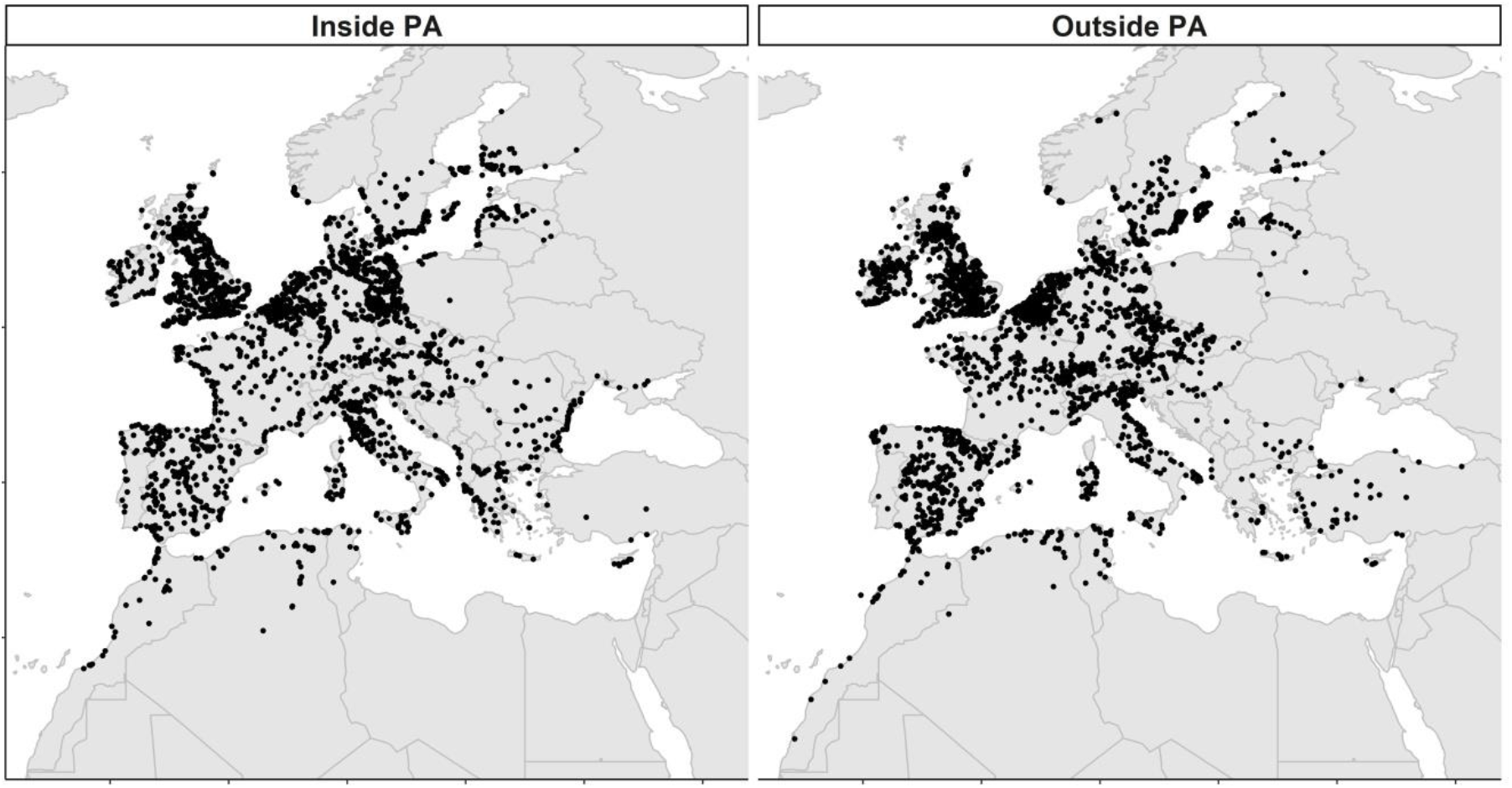
Map of the study area including 7,071 monitoring sites inside a protected area (PA, n = 3,374) and outside (n = 3,697), in 39 Western-Palearctic countries.

### Protected areas and temperature data

Site protection was reported for 3,374 sites from the World Database on Protected Areas (IUCN, UNEP-WCMC 2019), the Natura 2000 and the CDDA databases (www.eea.Europa.eu) (Fig. 2). Sites were considered as protected when their coordinates were included in the polygon of a protected area designated before 2017. When polygon data were absent (12% of the cases), a circular buffer was created based on the PA size reported in the World Database on Protected Areas (note that 100% concordance of site protection status was found by creating a circular area on the subset of PAs with polygons). The sites inside (n = 3,374) and outside (n = 3,697) PAs had a similar number of counts (in average (±SD) 16.8±5.7 and 16.4±5.7, respectively) and a similar spatial distribution (in average (±SD) Lat. 49.8±6.2, Lon. 7.0±9.1 and Lat. 50.3±6.1, Lon. 5.2±9.0, respectively, Fig. 2).

The HadCRUT4 dataset (Morice et al. 2012) that has a spatial resolution of 0.5° was our source of temperature data. Yearly winter temperatures were computed each winter as the average of the mean monthly temperatures of November, December and January.

### Community temperature indices

Winter species temperature indices (STI) were computed as the species thermal affinity across the non-breeding species distribution following Gaget et al. (2018) (adapted for non-breeding waterbirds from Devictor et al. (2008)). The winter STI is the long-term average January temperature (WorldClim database, 1950-2000, http://worldclim.org/) experienced by the species across the non-breeding (overwintering) distribution (extracted from www.birdlife.org 2015). Sub-species with distributions in Sub-Saharan African were removed to avoid an overestimation of the temperature experienced by the studied populations (Appendix 3).

The CTI and CTI standard deviation (CTI_sd_) were computed following Devictor et al. (2008) and Gaüzère et al. 2019 on species occurrence (presence/absence). The CTI is the average STI of the species present in the community per count event (see Appendix 4). The CTI_sd_ is the standard deviation of the species STI present in the community per count event, quantifying the STI heterogeneity in the community. Thus, the CTI increases over the years when there are more warm-dwelling species or fewer cold-dwelling species. The CTI_sd_ increases over the years when the thermal affinities of the community become more heterogeneous (Fig. 1). Occurrence data were used instead of abundance data to make it easier to interpret the processes of colonization-extinction, but usually produce similar CTI trends (e.g. Devictor et al. 2008, Gaget et al. 2018).

## Data analysis

### Protected areas, CTI, CTI_sd_ and climatic debt

Temporal changes of temperature, CTI and CTI_sd_ depending of the PA status were assessed with generalized linear mixed effects models (GLMM, Gaussian error distribution). The explanatory terms were the *year* (continuous variable from 1993-2017), the site *protection status* (Inside or Outside) and the interaction *year* × *protected status*. The *site identity* was added as a random effect on the intercept in the CTI and CTI_sd_ models. The spatial autocorrelation was taken into account by including the site geographical coordinates as an exponential spatial correlation structure in the model (Gaget et al. 2018). The linear model was:

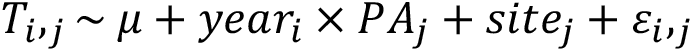

where *T*_*ij*_ was the temperature, CTI, or CTI_sd_, in year *i* at site *j*, μ was the intercept, PA was the site protection status of site *j*, site was the random intercept per site that follows a Gaussian distribution with mean of zero and variance σ^2^, and ε was the residual variance for each observation under a Gaussian distribution and an exponential spatial correlation. In order to visually assess whether it was appropriate to model inter-annual changes as a linear effect, we generated and plotted mean annual values (± 95% CI) by using the same model, but changing year to a categorical variable.

We looked for evidence of climatic debt accumulated by the waterbird communities by assessing the difference between the linear trends of temperature and CTI, following Devictor et al. (2008). First we investigated the latitudinal gradients in temperature and CTI with a GLMM (Gaussian error distribution), using the latitude as a fixed effect. The latitudinal gradient was converted to kilometres by dividing it by 111.128, i.e., the average number of kilometres per 1 decimal degree temperature over the whole study area. Then the temporal temperature change (°C yr^−1^) was converted to a velocity of temperature change in kilometres (km yr^−1^) by using the latitudinal temperature gradient (°C km^−1^) from South to North of the study area. The same was done with the CTI. Lastly, the climatic debt was obtained by subtracting the velocity of the CTI change from the velocity of the temperature change over the study period.

In addition, we assessed the temporal trend of cold- and warm-dwelling species inside *vs.* outside PAs. Species were classified in two categories as ‘cold-dwelling’ or ‘warm-dwelling’ following their STI in relation to the individual site, i.e., cooler or warmer than the mean CTI of the site’s time series, respectively. Then, the number of cold and warm-species was summed per survey. We used these two simplified categories to control the accuracy of the community thermal changes assessed with CTI and CTI_sd_. The temporal changes in number of cold- and warm-dwelling species were assessed using in a GLMM (Poisson error distribution) with fixed effects of year, the thermal-dwelling category (cold or warm), the site PA status (Inside or Outside) and their three-way interactions. The site identity was added as a random factor. The spatial autocorrelation was taken into account by including the site geographical coordinates as an exponential spatial correlation structure in the model.

### Community changes in response to protected area surface

We investigated whether the local CTI, climatic debt and CTI_sd_ trends were correlated with the local PA surface. First, a moving-window approach was used to investigate the spatio-temporal changes of temperature, CTI, climatic debt and CTI_sd_. The study area was divided in 1,032 cells of 5°×5° resolution (c. 500×500 km) spaced from each other by one latitudinal or longitudinal degree. We performed one GLMM per cell per response variable (temperature, CTI and CTI_sd_), to investigated their change over years using the same model structure as before. Temperature, CTI and climatic debt spatio-temporal changes were given in km yr^−1^, and in °C yr^−1^ for the CTI_sd_. Note that each cell included both protected and not protected sites and at least 15 sites (mean of 175 sites), to avoid cells with a small number of sites at the edge of the study area.

Then, we investigated the relationship between the PA surface per cell and the CTI spatial shift, CTI_sd_ and climatic debt trends, estimated from the models above. One generalised linear model (GLM, Gaussian error distribution) was used per response variable with fixed effects the PA surface (sum of the PA surfaces per cell, log transformation assuming a non-linear relation) and the temperature spatial shift plus their interaction to control for the climate warming pressure. To investigate the geographical PA surface repartition in the Western-Palearctic, we also assessed in a GLM whether PA surface increased with latitude, longitude and their interaction.

All the statistical analyses were performed with R 3.4.3 (R Core Team 2017), using the ‘glmmTMB’ package (Magnusson et al. 2017) for the GLMM and GLM.

## Results

### Protected areas, CTI, CTI_sd_ and climatic debt

The temperature increased by 0.04°C per year (P < 0.001) without significant difference between inside and outside PA (P = 0.2, Table 1, Fig. 3a). The CTI increased faster inside PAs than outside, about 0.010°C yr^−1^ to 0.006°C yr^−1^, respectively (Table 1, Fig. 3c). CTI_sd_ increase was significant inside PAs, but not significant outside PAs (Table 1, Fig. 3d). Therefore, within PAs, the results matched scenario 3 (Fig. 1; colonization only), whereas outside PAs the results matched scenario 4 (Fig. 1; colonization and extinction).

**Table 1:**
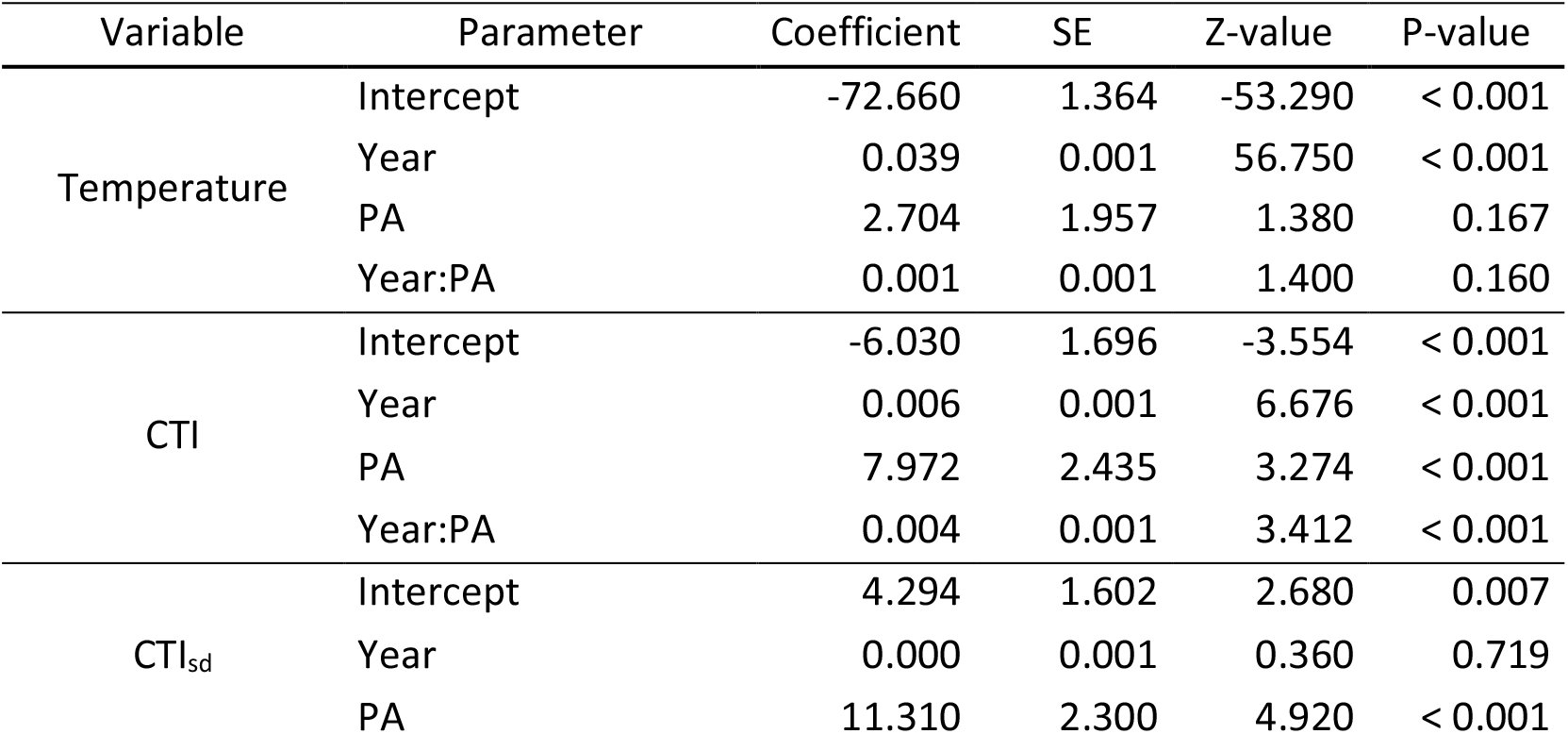
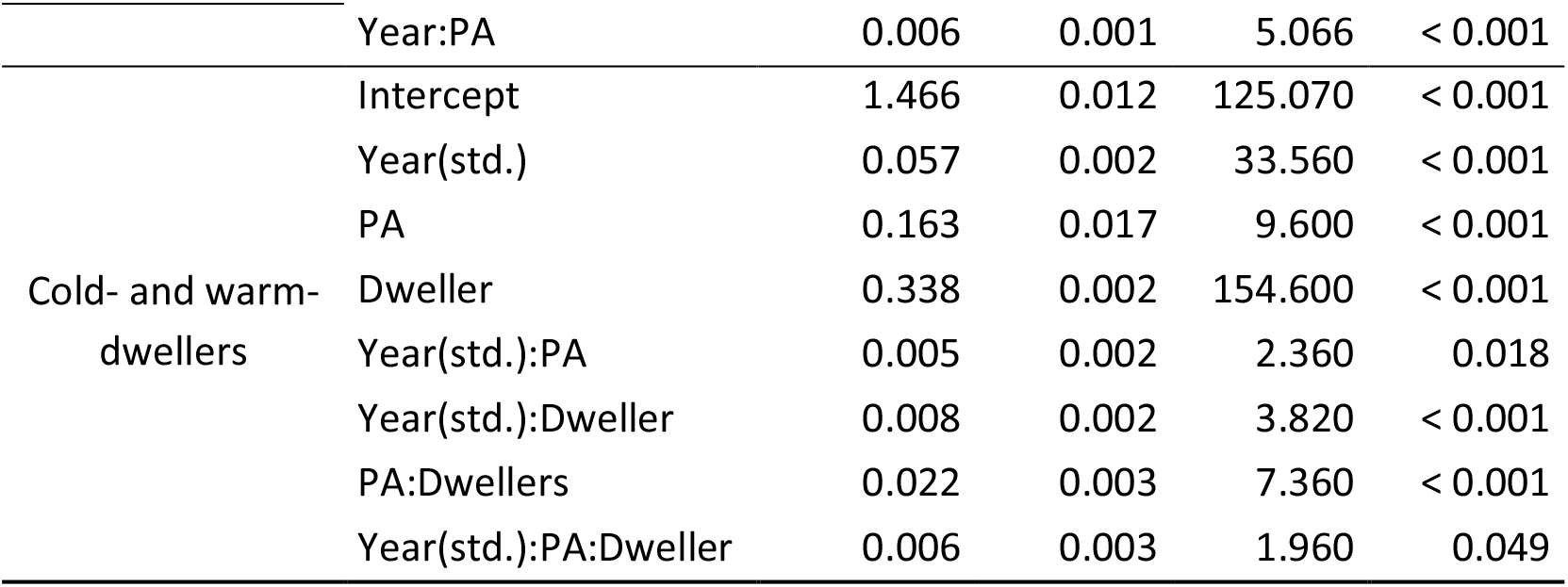
Temporal trends of the temperatures, community temperature index (CTI) and standard deviation of the CTI (CTI_sd_) and number of cold- and warm-dwelling species regarding the protected area (PA) site status. Base line is sites outside PA and cold-dwelling species. Years were standardized to zero mean (std.) in the thermal-dwellers model and interactions are notified by ‘:’.

**Figure 3:**
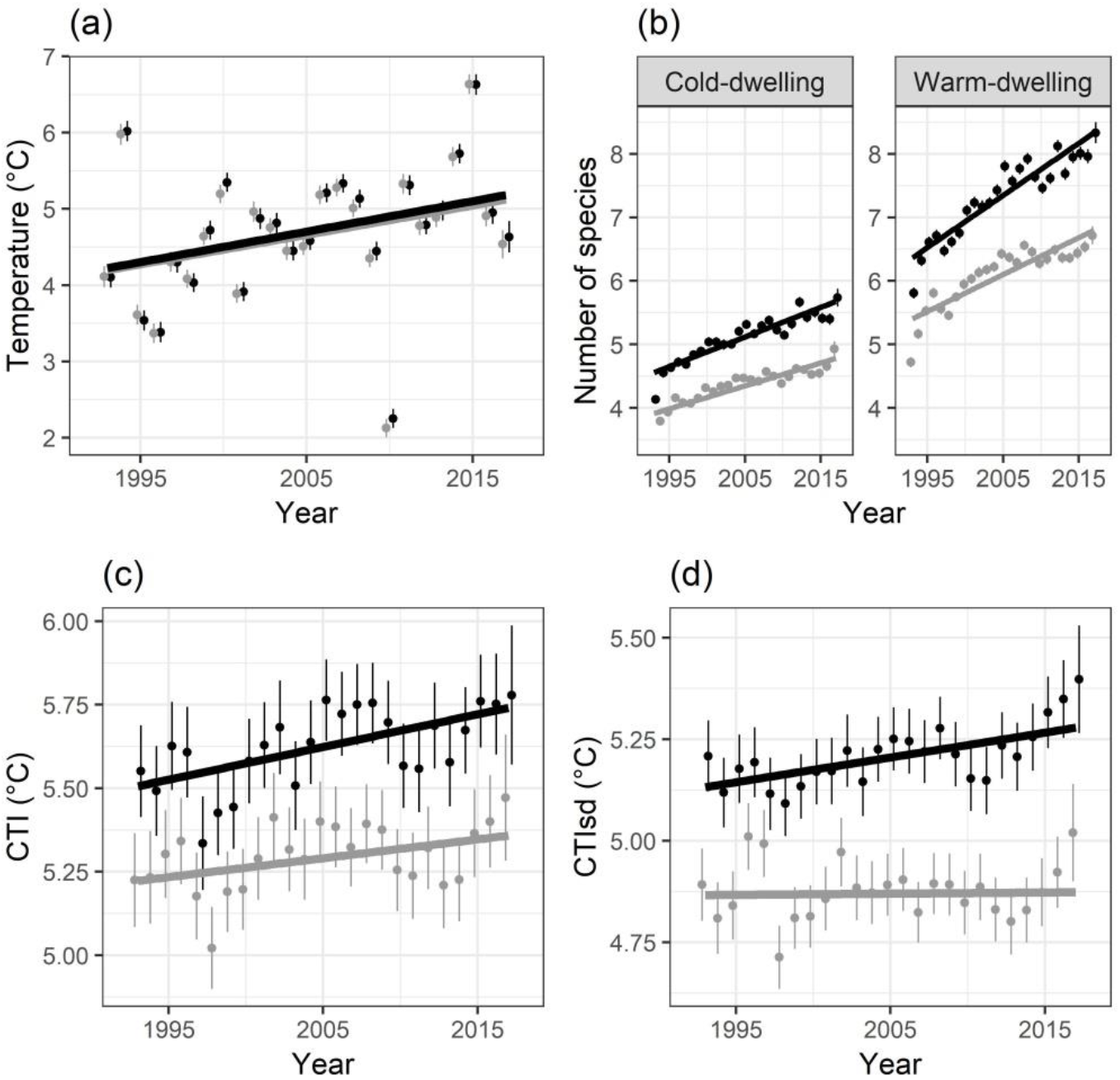
Temporal trends inside PA (black) and outside PA (grey) of the (a) temperature, (b) number of cold- and warm-dwelling species, (c) community temperature index (CTI) and (d) standard deviation of the CTI (CTI_sd_). Mean (± 95% CI) are represented by points.

Temporal changes in CTI lagged behind changes in temperature. The temperature latitudinal gradient was about −0.38°C per decimal degree (SE = 0.005, Z = −78.75, P < 0.001) and −0.31°C (SE = 0.004, Z = −69.56, P < 0.001) for the CTI. The temperature increase was equivalent to a latitudinal shift of 11.4 km yr^−1^ (285 km in 25 years). The temporal CTI trend was equivalent to a shift 43% larger inside PAs than outside, with about 3.5 km yr^−1^ inside the PAs (87 km over 25 years) and 2.0 km yr^−1^ outside (50 km over 25 years). Consequently, the climatic debt was about 7.9 km yr^−1^ inside the PAs and 9.4 km yr^−1^ outside (198 km and 235 km over 25 years, respectively).

The number of cold- and warm-dwelling species both increased significantly over the study period, but the trends and average numbers of species were significantly greater inside PAs (Table 1, Fig. 3b). Warm-dwelling species were more numerous and their number increased faster than the cold-dwelling species (Table 1). Inside PAs, the warm-dwelling species increased also faster than the cold-dwelling species (Table 1). This suggests that, both inside and outside PAs fit between scenarios 3 and 4 – with more colonization than extinction.

#### Community changes in response to protected area surface

The temperature increased significantly in 80% of the study area, with the exception of the northern half of the Iberian Peninsula (Fig. 4A). The CTI significantly increased in 37% of the cells (384/1,032), mostly from South Balkans to West France and around the Baltic Sea (Fig. 4B). Consequently, there was climatic debt in 66% of the area, mostly in the northern half of Europe (Fig. 4C). Lastly, the CTI_sd_ trend was significantly positive in 39% of the cells, mainly in the East and the South, but also around the Baltic Sea (Fig. 4D).

**Figure 4:**
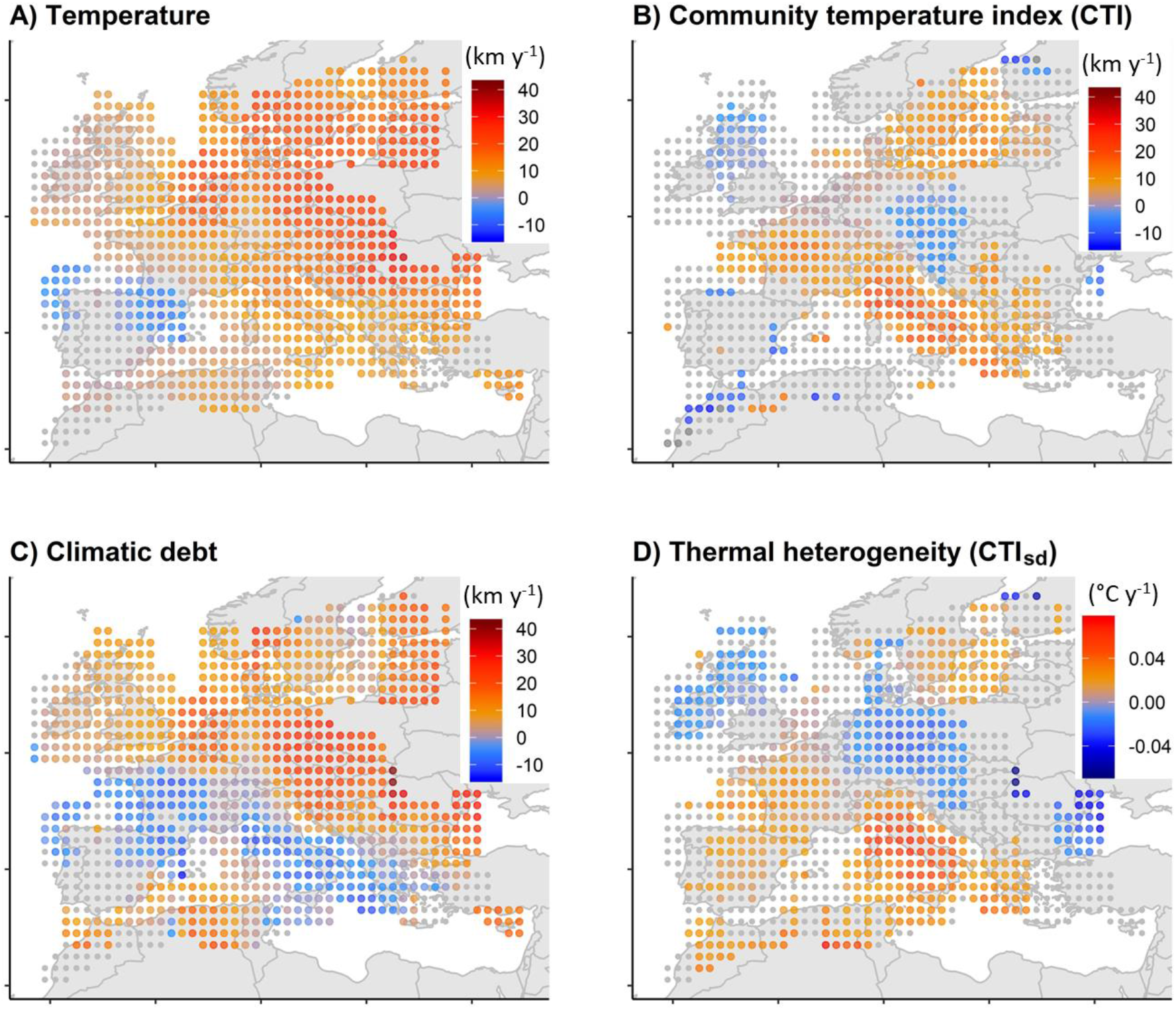
Spatio-temporal trends from 1993-2017 of (A) temperature, (B) community temperature index CTI, (C) climatic debt and (D) thermal heterogeneity CTI_sd_. Trends are represented by points located at the centre of the corresponding cell (5°×5° resolution). Coloured points denote a significant trend (P < 0.05, positive ‘red’, negative ‘blue’) while colour gradient indicates the trend intensity (not significant trend in grey).

The CTI spatial shift increased with PA surface and temperature spatial shift (P ≤ 0.001) but without a significant interaction (Table 2). Consequently, the climatic debts accumulated were smaller where there was more PA surface and greater where the temperature spatial shift was faster (P ≤ 0.001) (Table 2). The temporal CTI_sd_ trends did not change with PA surface (P = 0.3), but increased less where the temperature spatial shift was faster (P < 0.001, Table 2). However, the CTI_sd_ trends decreased when both PA surface and temperature spatial shift increased as demonstrated by a negative interaction between temperature spatial shift and PA surface (Table 2). The PA surface areas were greater in southwest and northeast, as the PA surface decreased with the longitude (β = −0.266, P < 0.001) but not with the latitude (β = −0.067, P < 0.14), with a positive and significant interaction (β = 0.274, P < 0.001, Appendix 5).

**Table 2:**
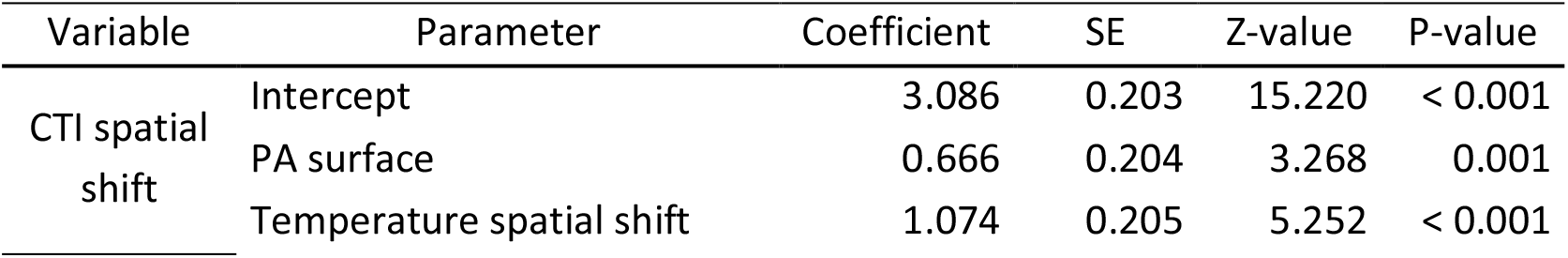
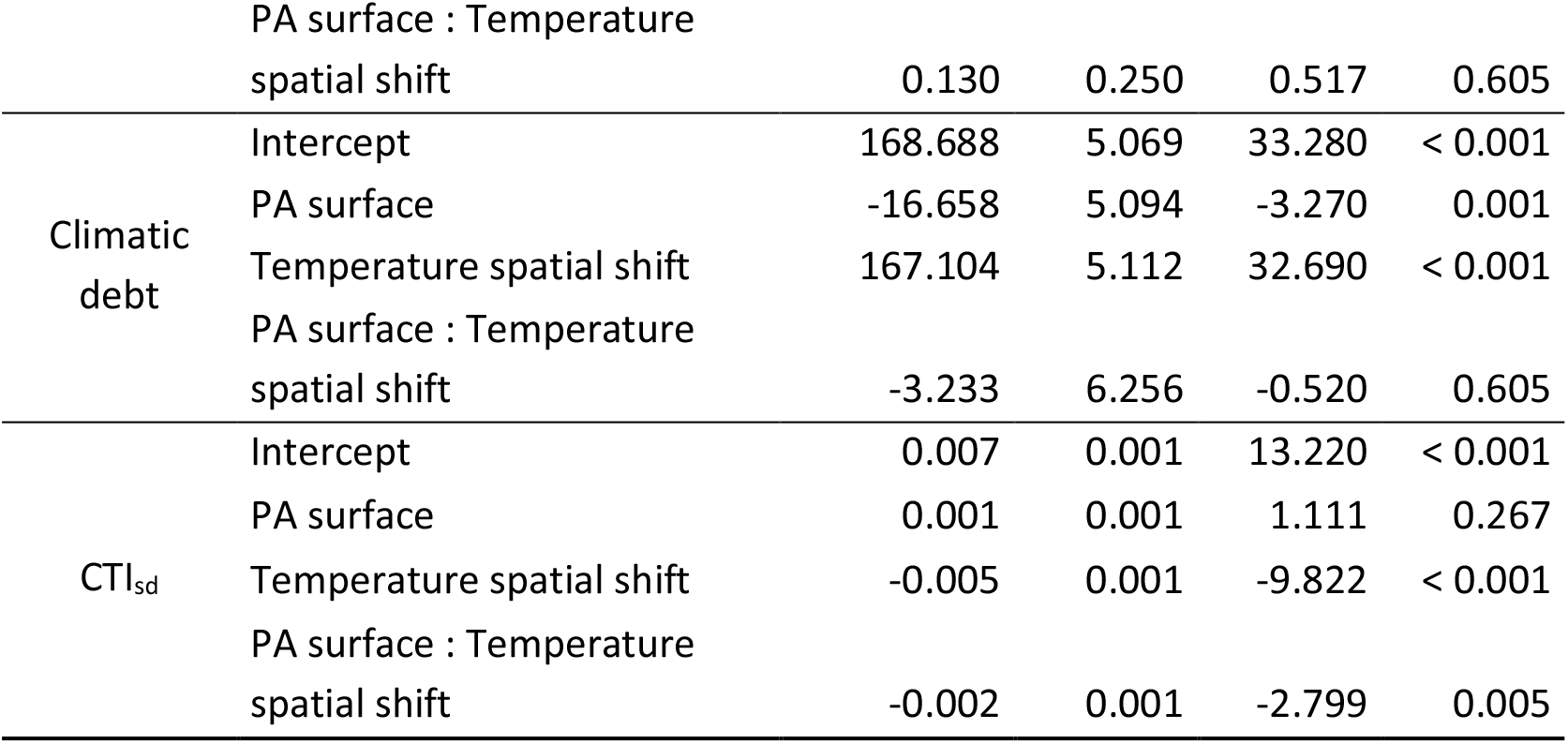
Spatial effect of protected area surface (log transformed) and interaction with the temperature spatial shift on the CTI spatial shift, climatic debt and CTI_sd_, per cell of 5°×5°. Interacting effects denoted by ‘:’.

## Discussion

### Community adjusts faster to climate warming inside protected areas

This study represents one of the first empirical and international assessments addressing difference in community changes in response to climate warming within PAs on a continental scale. We found that the CTI faster increase inside PAs compared to outside areas was driven mainly by colonization from warm-dwelling species, which is consistent with other studies on birds and other taxonomic groups (Thomas et al. 2012, Gillingham et al. 2015). Indeed, when looking at finer spatial scale, the CTI increase was more positive where PA surfaces were larger, suggesting a positive relationship of PA coverage on community thermal changes (Gaüzère et al. 2016).

Overall, we find that non-breeding waterbirds in the Western-Palearctic show a climatic debt, but this debt is 16% lower inside PAs. Communities inside PAs had higher colonization, lower extinction and lower climatic debt. Moreover, PAs supported higher waterbird species richness, which is consistent with the PA designation on wetlands of high biological importance, e.g. by the Ramsar Convention and the European Union’s Nature Directives. Therefore PAs are not only important to reduce the direct anthropic pressures (Godet & Devictor 2018) but also are associated with reduced climatic debt. Such conservation benefit was expected by international conservation policies (Trouwborst 2009, 2011, 2012), which use PAs and species protection status as the main conservation measures to buffer the negative impacts of climate change, in order to reduce ecosystem pressures and promote species adaptation to climate change (Trouwborst 2011, 2012). The Western-Palearctic falls under several of these international conventions, such as the Ramsar, Bern and Bonn Conventions, and the benefits provided by habitat and species protection (Gamero et al. 2017, Pavón-Jordán in Rev.) seem effective to facilitate the species adjustment to climate warming (Gaget et al. 2018). For example, both the Great Cormorant (*Phalacrocorax carbo*) and the Great Egret (*Ardea alba*) had declining populations in Europe until their designation as protected species by the Bern Convention (19.IX.1979) and the Birds Directive (79/409/EEC) in 1979. After that, a fast population recovery occurred notably by a northward expansion (Hiley et al. 2013, Ławicki 2014, Marion & Bergerot 2018).

Species richness of non-breeding waterbird increased over the study area, particularly inside PAs, in line with recent general positive trends of Western-Palearctic waterbird populations (Amano et al. 2018). Furthermore, inside – but not outside – PAs the variation in CTI (CTI_sd_) increased over time, and we find a general increase in CTI of both cold- and warm-dwelling species over time. These findings suggest that inside PAs, species with high thermal affinity colonized the community, but at the same time species with low thermal affinity were less likely to be locally extinct. In other words, PAs can act as refuge by improving species resilience again climate warming (Santangeli et al. 2017, Berteaux et al. 2018), likely by ensuring ecological requirements needed for species persistence despite the proximity with the thermal niche edge.

### Heterogeneity of temperature and community changes

The intensity of the winter temperature warming increased over a southwest-northeast gradient, driving the community adjustment through a similar gradient of intensity, although not perfectly (Fig. 4). The thermal isocline shift towards the northeast is related to the continental shape and the oceanic influence of the Gulf Stream (IPCC 2014). Interestingly, the non-significant temperature and CTI trends in the southwest of the Western-Palearctic resulted in negligible climatic debts. Conversely, the climatic debt increased in the northeastern countries where strong temperature warming occurred (Fig. 4), which non-breeding waterbirds were not able to fully track.

Temperature was likely not the only aspect of the physical environment that constrained species’ distributions. The local pattern of CTI changes contrasted with the expected relative increase of warm-dwelling species. While several other factors are likely to have affected species’ distributions, the CTI focuses on species assemblage changes in response to temperature changes, but its trend can also be affected by other drivers of population change (Bowler & Böhning-Gaese 2017). For example, in the UK, despite a species-specific west-east waterbird redistribution (Austin & Rehfisch 2005), the CTI changes were likely altered by the recent increase of geese and the decrease of waders (Frost et al. 2019), which have low and high STIs, respectively (Appendix 3). Consequently, the subsequent community reshuffling may jeopardize the detection of a community thermal adjustment, if it exists (Bowler & Böhning-Gaese 2017). Similarly, the absence of CTI increase in Central Europe and in the Netherlands despite the temperature increase should encourage species-specific investigations (e.g. Pavón-Jordán et al. 2015). Such unexpected population changes, under the hypothesis of a community adjustment to climate warming, increase the theoretical mismatch between community and temperature changes (Kerbiriou et al. 2009, Galewski and Devictor 2016).

Although milder climate conditions reduce ice and snow in the northern regions and enhance northward range expansion (Brommer 2008, Schummer et al. 2010), the community adjustment to climate warming was not particularly strong in northern Europe (Fig. 4). This may be the result of average temperatures not accurately reflecting the thermal conditions that affect species’ distribution. For example, in the northern regions, severe cold spells may cause potential large mortality, this limiting species distribution changes (Gunnarsson et al. 2012).

Considering the strong waterbird distribution change in northern Europe (Brommer 2008, Lehikoinen et al. 2013), the lack of CTI increase also suggests some limits of the CTI framework. The CTI measures changes of species assemblages (Devictor et al. 2008) and could be sensitive to the number of species already present in the community. Indeed, when there are few species at the beginning of the monitoring, because of ice cover for example, the CTI trend should be more sensitive to the species arrivals. We acknowledge that we didn’t tack into account for this potential uncertainty. Consequently, our ability to measure species distribution change is challenged in these ice-dominated regions, where the community adjustment to climate warming is likely underestimated (Fox et al. 2019).

### Perspectives for research and conservation

Indicators are essential tools to synthesize population dynamics and inform public policies (Tittensor et al. 2014). The CTI is an intuitive indicator with which to measure and communicate the impact of climate warming on communities (Devictor et al. 2012, Gaüzère et al. 2019). Here, we go one step further and used the CTI_sd_ to identify the colonization-extinction patterns in response to climate warming (see also Appendix 1). With these simple indicators, we identified that the community adjustment to temperature was mainly due to colonization by the warm-dwelling species inside PAs, while outside PAs the extinction of the most cold-dwelling species was nearly equivalent to the colonization by warm-dwelling species (Fig. 3d).

This study relied on an international coordinated monitoring program, allowing us to investigate whether community adjustment to climate warming was higher in PAs. The IWC is a monitoring scheme that aims to ensure waterbird counts (full check-lists) in both protected and unprotected areas (Delany 2010). However, we acknowledge that PAs were not randomly distributed (Fig. 2), and that such non-randomness could induce spatial aggregation between PA density and CTI changes. Nevertheless, when looking at the spatio-temporal changes (Fig. 4), spatial aggregation was moderate. In particular, the CTI trends were heterogeneous even between areas with high PA density (Fig. 2), e.g. in the Netherlands or southern UK. More emphasis should be given to investigate how PA characteristics, e.g. management plans, influence at a local scale community adjustment to climate warming (Monzón et al. 2011).

Non-breeding waterbirds have high capacity to respond to climate warming by a distribution change (Maclean et al. 2008, Lehikoinen et al. 2013, Pavón-Jordán et al. 2019) even more than other groups of birds (Brommer 2008). Our study reveals a faster average distribution shift, 2.0 to 3.5 km yr^−1^, in comparison to the European common breeding birds (2.1 km yr^−1^, Devictor et al. 2012) and other taxa (1.8 km yr^−1^, Chen et al. 2011). Indeed, most of the Western-Palearctic waterbirds are migratory, spending energy and facing multiple threats during migration. Shortening their migration routes, by overwintering at more northern latitudes, could be advantageous, by decreasing the migration cost and benefits their fitness (Gilroy 2017, Reneerkens et al. 2019).

These rapid distributional changes that we found bring into question the future effectiveness of the PA networks, because of the locations of these sites potentially do not match the future distributions of waterbird species (Araújo et al. 2004). In the Western-Palearctic, the PA network covers 45% of the inland wetlands (Bastin et al. 2019) and even if the number of PAs increases in the northeast, the network still does not cover all the wetlands important for waterbird conservation (Pavón-Jordán et al. 2015, Pavón-Jordán in Rev.). In the future, PAs can maintain their conservation value if the extinction of conservation concern species is compensated by the colonization of other species of conservation concern (Hole et al. 2009, Johnston et al. 2013
). From that perspective, our results are encouraging, as they indicate the PAs would still remain important for waterbird conservation in the future, since waterbird colonization was greater, and extinction lower, inside a PA compared to outside. However, more studies are needed to understand the mechanisms by which PAs have buffered against climate change, and to evaluate the current and future completeness of the PA network particularly for conservation concern species (Pavón-Jordán et al. 2015).

## Acknowledgements

We acknowledge the thousands of volunteers and professionals involved in the International Waterbird Census, making this research possible. This research was funded through the 2017-2018 Belmont Forum and BiodivERsA joint call for research proposals, under the BiodivScen ERA-Net COFUND programme, with the following funding organizations: the Academy of Finland (Univ. Turku: 326327, Univ. Helsinki: 326338), the Swedish Research Council (Swedish Univ. Agric. Sci: 2018-02440, Lund Univ.: 2018-02441), the Research Council of Norway (Norwegian Instit. for Nature Res., 295767), the National Science Foundation (Cornell Univ., ICER-1927646) and we also acknowledge the Swedish Environmental Protection Agency.

## Appendix 1. Colonization and extinction patterns revealed by the CTI_sd_

## 1. Simulations of the species extinction/colonization in response to temperature increase and subsequent changes of Community Temperature Index (CTI) and standard deviation (CTI_sd_) over time

Following the Figure 1, four scenarios were simulated (Rcode below). The scenarios were: (1) ‘No colonization-No extinction’; (2) ‘Extinction only’; (3) ‘Colonization only’; (4) ‘Colonization-Extinction’.

For each of the four scenarios, we simulated an occurrence matrix for 100 species considered in three temperature dwelling classes over 25 years (from 1 to 25) and 100 sites. We attributed different Species Temperature Index (STI) values to the species from a random simulation of STI values based on a Gaussian distribution of mean 0 and SD 10. Twenty five species were considered as extreme cold-dwelling species with STI inferior to −5°C, 50 species were considered as slight cold- or warm-dwelling species with STI between −5°C and 5°C and 25 species were considered as extreme warm dwelling species with STI superior to 5°C. Species occurrence were simulated from a binomial distribution with different probabilities between the extreme cold-dwelling (p=0.25 or p=0.25-year/100, if extinction), slight cold- or warm-dwelling species (p=0.75) and extreme warm-dwelling species (p=0 or p=year/100, if colonization). From the 100 original species pool, 1 to 90 species were randomly removed in order to simulated different environmental filters. We computed the CTI and CTI_sd_ values per year per site (see Methods). We used generalized linear mixed effects models (GLMM, Gaussian error distribution) with the CTI or CTIsd as the response variable, the year as the explanatory term and the site in random effect. Finally, the estimate temporal slope and its p-value were collected. We simulated the four scenarios 100 times following this process (Rcode below).

Rcode used for the simulations:

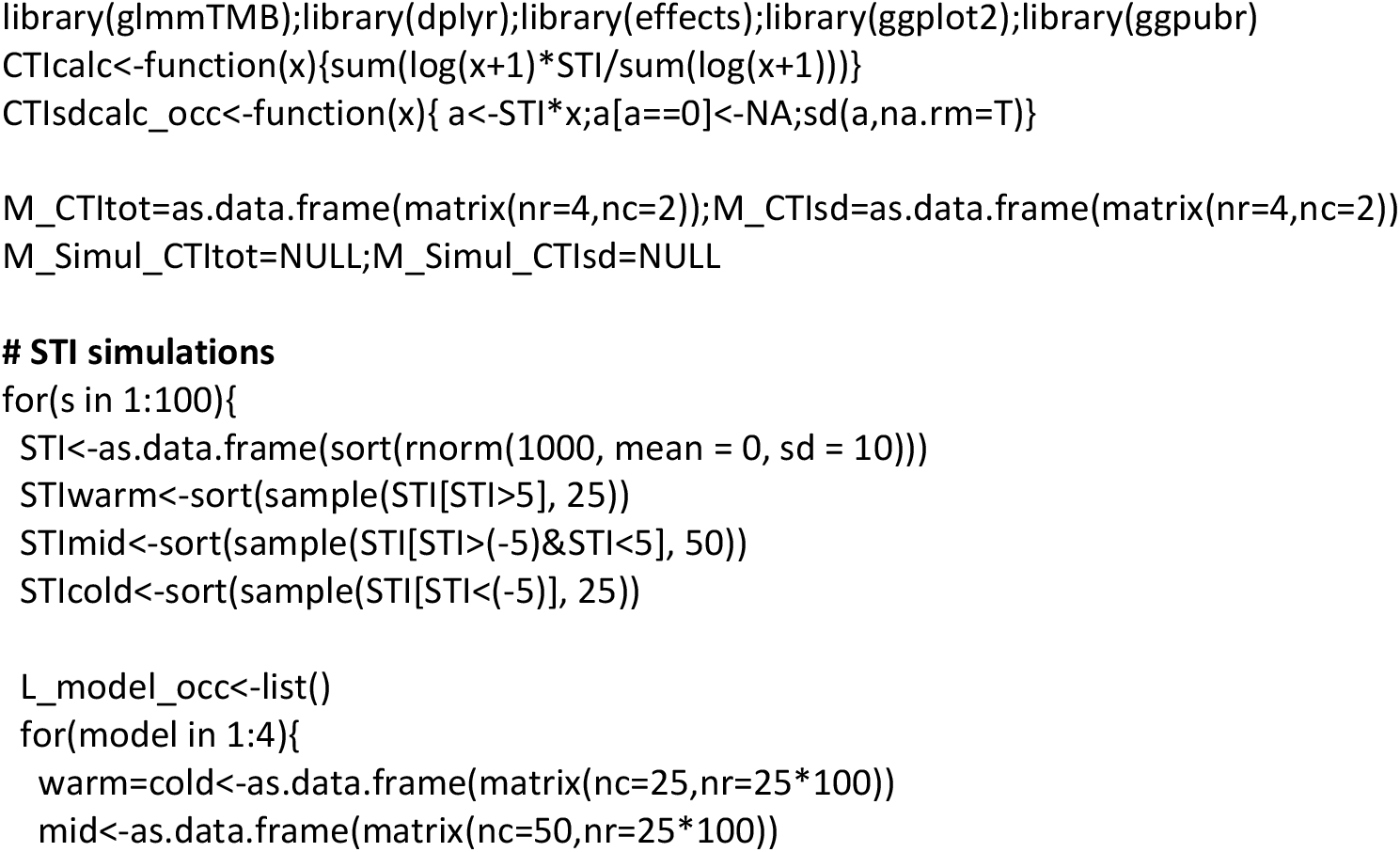
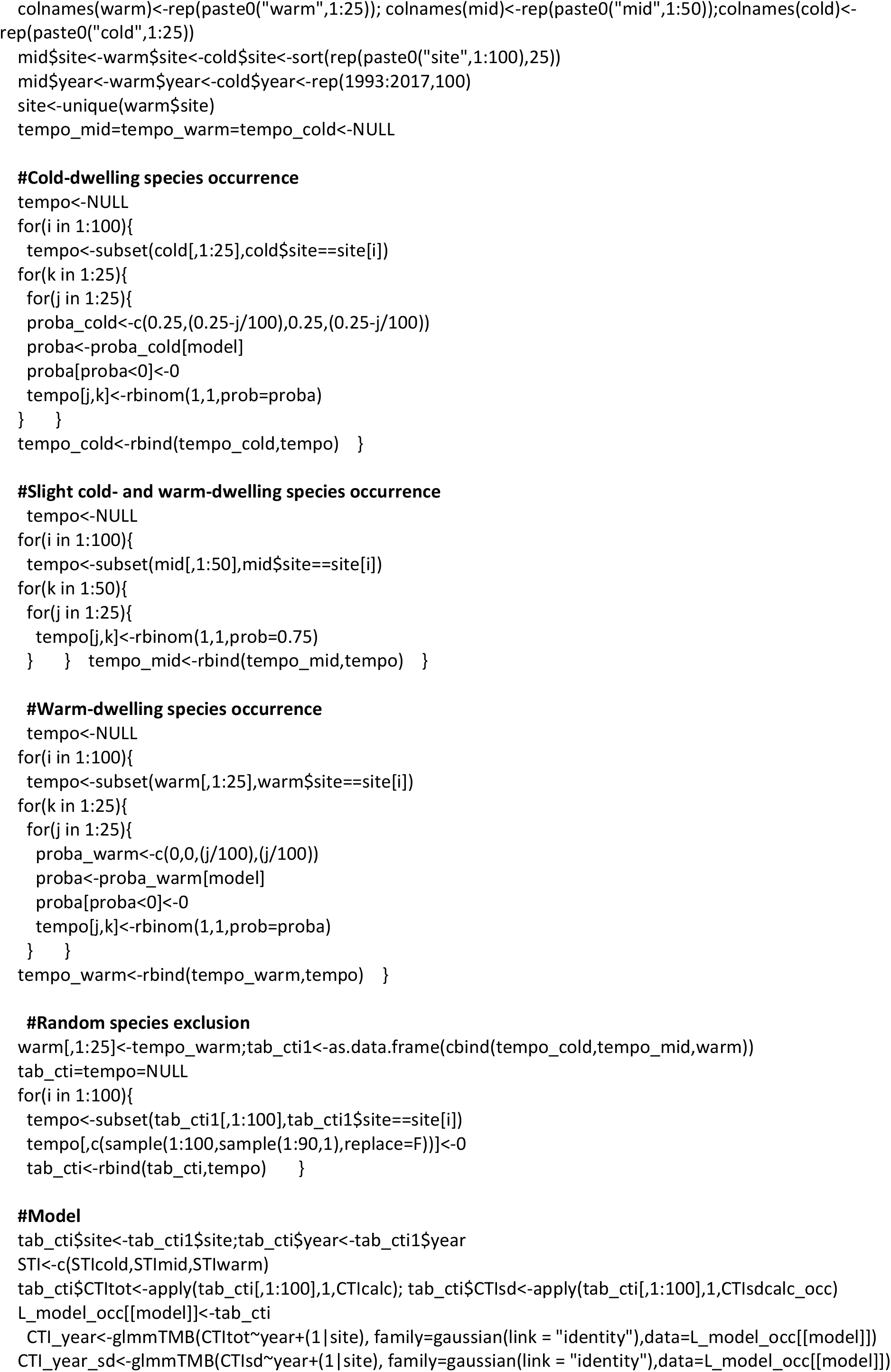
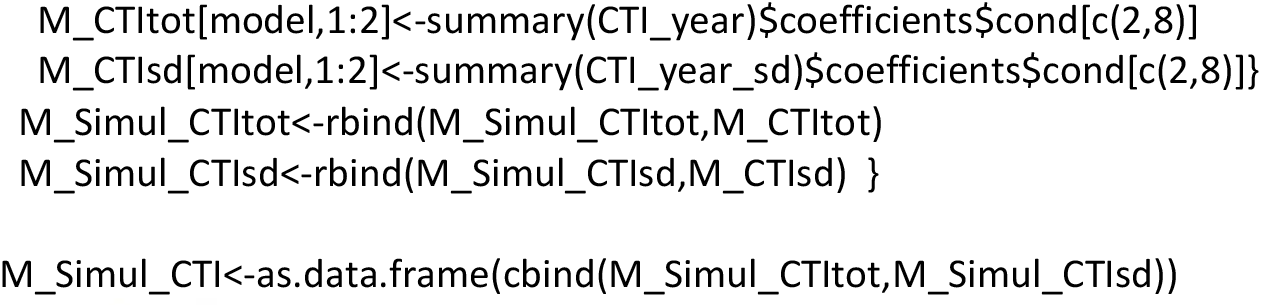

**Figure S1a:**
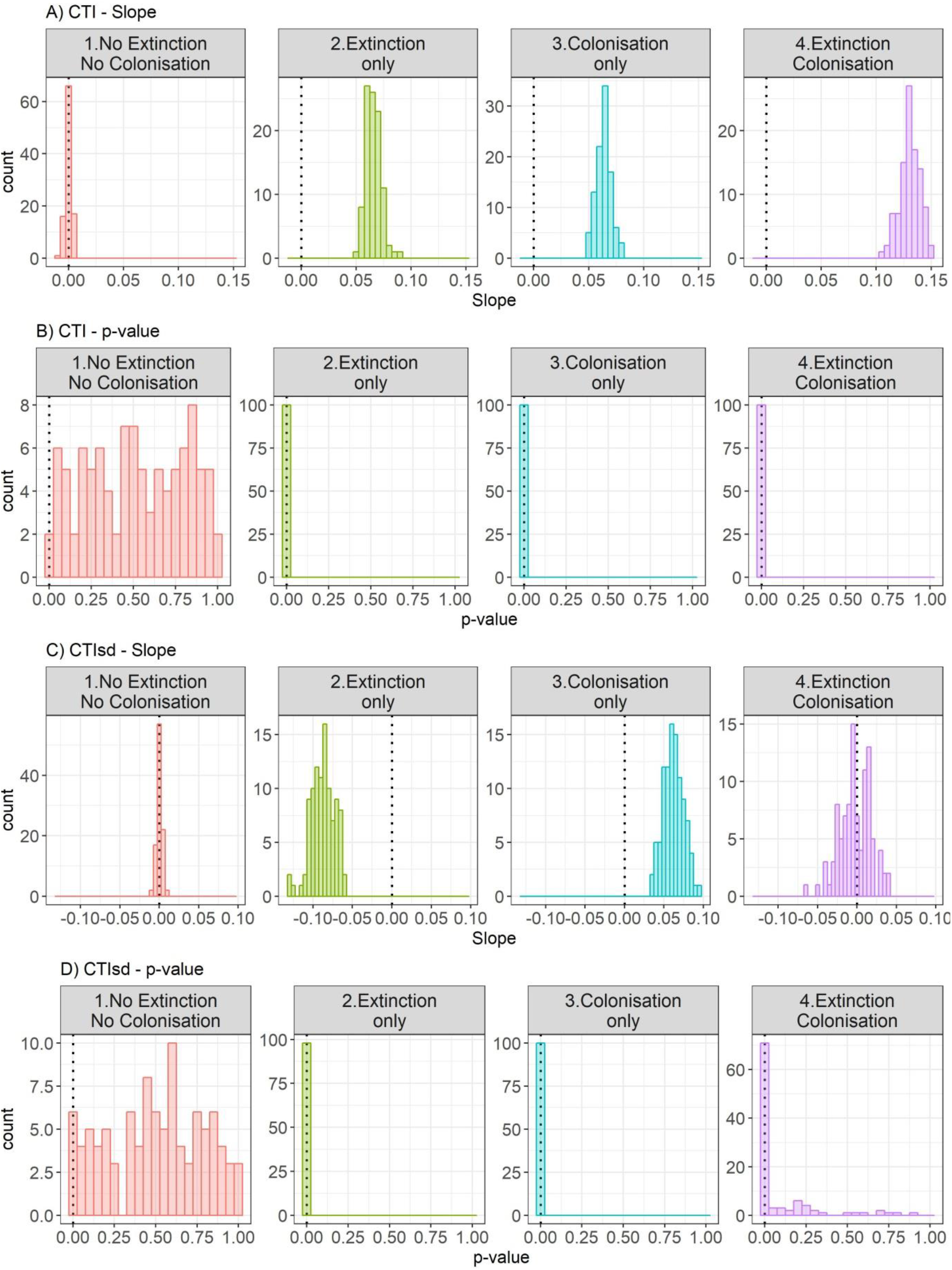
Histograms of the model outputs per scenario. A) CTI estimated slope, B) p-value corresponding to the CTI slope, C) CTIsd estimated slope, D p-value corresponding to the CTIsd slope. The scenarios of community changes in response to temperature increase were: (1) ‘No colonization-No extinction’; (2) ‘Extinction only’; (3) ‘Colonization only’; (4) ‘Colonization-Extinction’.

## 2. Empirical observation of waterbird species extinction/colonization in response to temperature increase and subsequent changes of Community Temperature Index average (CTI) and standard deviation (CTI_sd_) over time

We highlighted the ability of the CTI_sd_ to be an indicator of colonization and extinction processes in response to climate warming. Indeed, community changes in response to temperature increase should result in four scenarios: (1) ‘No colonization-No extinction’ causes no CTI and CTI_sd_ changes; (2) ‘Extinction only’ causes CTI increase and CTI_sd_ decrease by the loss of cold-dwelling species; (3) ‘Colonization only’ causes CTI and CTI_sd_ increase by the gain of warm-dwelling species; (4) ‘Colonization-Extinction’ causes CTI increase by the species thermal turn-over, but no CTI_sd_ directional change (Fig. 1). We classified the count events in the four scenarios of colonization and/or extinction events, following what happening between the monitoring year and the next one (e.g., if between the counts *i* and *i+1* only one species colonized the site, the count *i* correspond to the scenario (3) ‘Colonization only’). For each count event, we measure the change of CTI_sd_ from a monitoring year and the next one (i.e., ΔCTI_sd_), which is supposed to be superior, inferior or equal to zero depending of the four colonization/extinction scenarios in response to temperature increase. We used a GLMM per scenarios (Gaussian error distribution) to investigate if the ΔCTI_sd_ values correspond to the expected patterns following the four scenarios. Site was added in random factors.

Conformely to the expectation under a community adjustment to climate warming, the ΔCTI_sd_ values was null in case of no extinction and no colonization (β = 0.000, P = 1), significantly negative in case of extinction only (β = −0.582, P < 0.001), significantly positives in case of colonization only (β = 0.590, P < 0.001), and not significantly different from zero in case of extinction and colonization (β = −0.005, P = 0.5) (Fig. S1b).

**Figure S1b:**
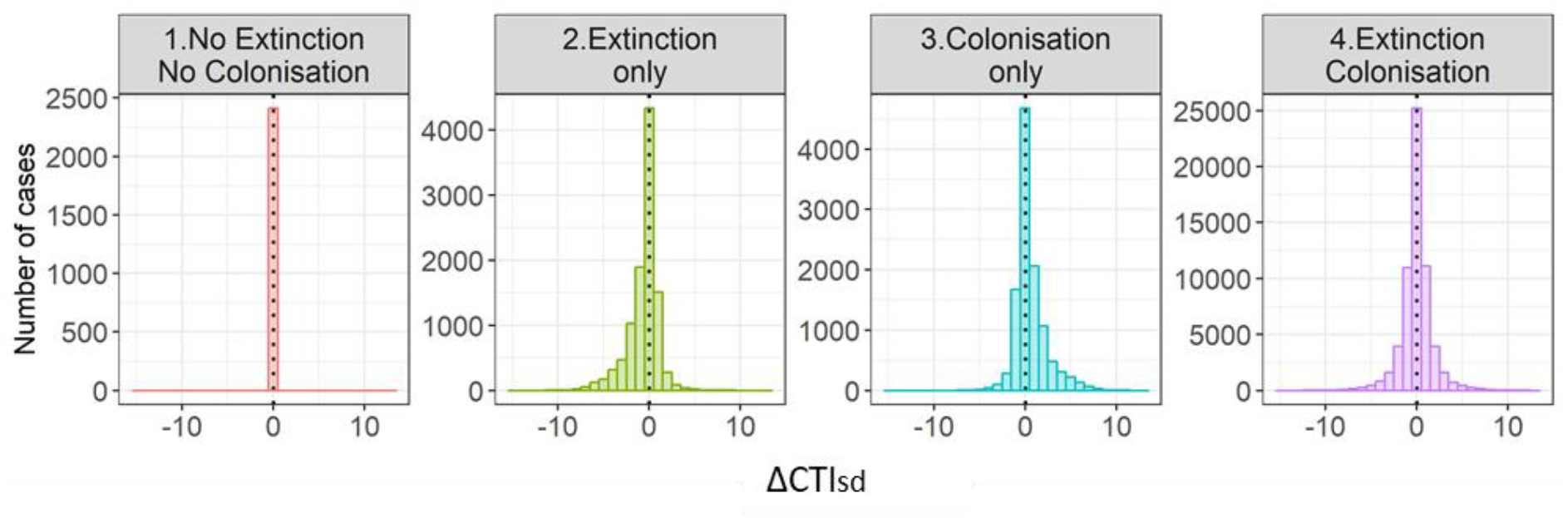
Histograms of the ΔCTI_sd_ values over the four scenarios of community changes in response to climate warming: (1) ‘No colonization-No extinction’ causes no CTI_sd_ changes, (2) ‘Extinction only’ causes CTI_sd_ decrease by the loss of cold-dwelling species, (3) ‘Colonization only’ causes CTI_sd_ increase by the gain of warm-dwelling species, (4) ‘Colonization-Extinction’ causes no CTI_sd_ directional change (Fig. 1). The dotted line is positioned on the zero to signify the absence of CTI_sd_ change.

## Appendix 2. Details of the monitoring per country

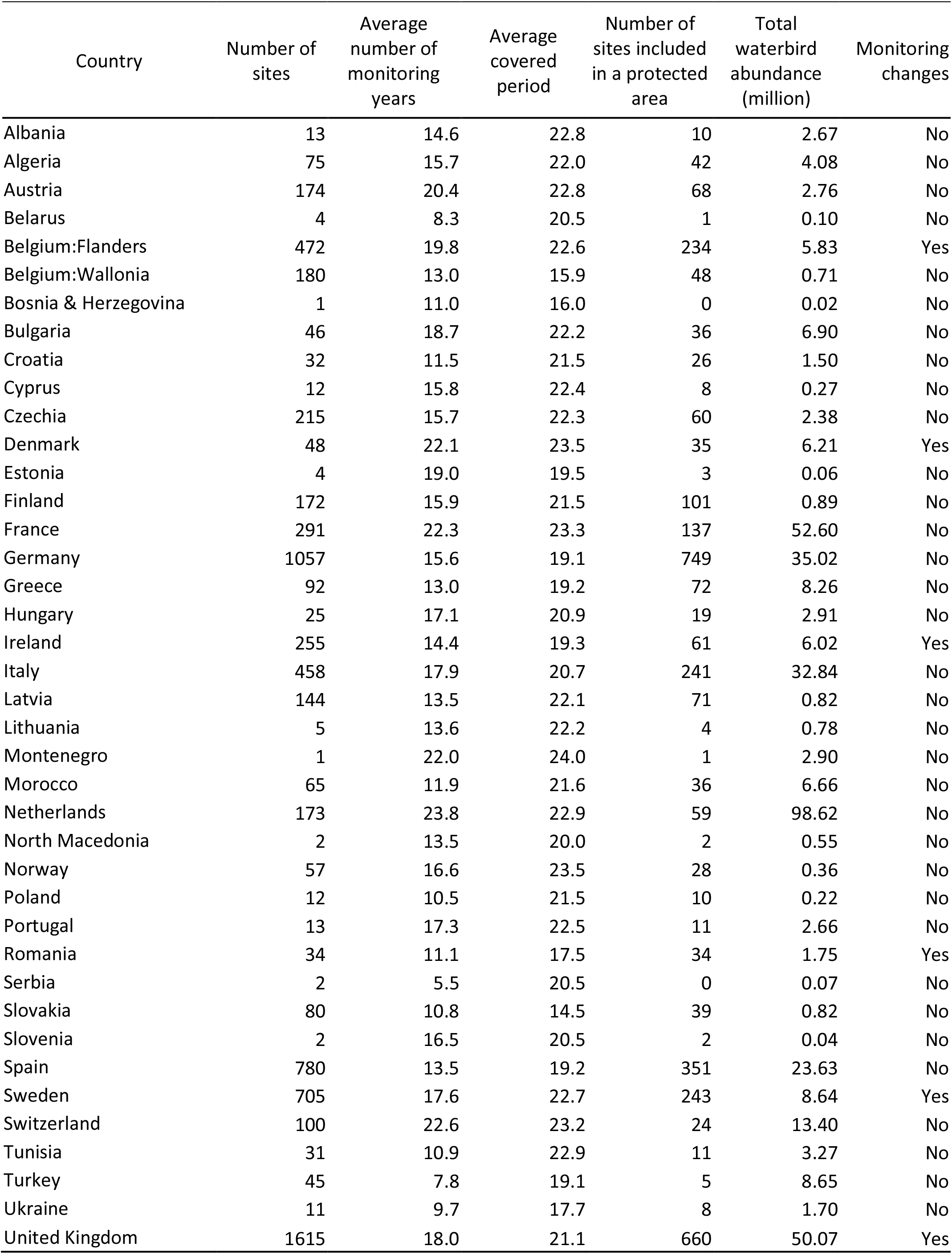

## Appendix 3. Additional species information

The winter STI is the long-term average January temperature (WorldClim database, 1950-2000, http://worldclim.org/) experimented by the species across its non-breeding (overwintering) distribution (extracted from www.birdlife.org 2015) only inside the African-Eurasian region defined by the African-Eurasian Migratory Waterbird Agreement (AEWA, http://www.unep-aewa.org). We removed the distribution of the sub-species resident in sub-Saharan Africa to avoid an overestimation of the thermal affinity tolerated by the studied populations (Involved species: *Ardea alba, Ardea cinerea, Botaurus stellaris, Gallinula chloropus, Phalacrocorax carbo, Podiceps cristatus, Podiceps nigricollis, Porphyrio porphyrio* and *Tachybaptus ruficollis*). Species considered as vagrant when their overwintering distribution was not included in the AEWA area and in the Western-Palearctic with a minimum threshold of 500 individuals over the 25 years.

**Table S2:** List of the species with their species temperature index (STI) and the number of sites occupied at least once.

| Scientific name | STI | Number of sites occupied |
| --- | --- | --- |
| *Actitis hypoleucos* | 23.16 | 1274 |
| *Anas acuta* | 16.90 | 2696 |
| *Anas clypeata* | 14.91 | 4929 |
| *Anas crecca* | 12.72 | 6854 |
| *Anas penelope* | 16.53 | 1642 |
| *Anas platyrhynchos* | -0.02 | 3770 |
| *Anas strepera* | 11.73 | 734 |
| *Anser albifrons* | 2.53 | 93 |
| *Anser anser* | 4.47 | 1229 |
| *Anser brachyrhynchus* | 2.02 | 2246 |
| *Anser erythropus* | 2.43 | 6239 |
| *Anser fabalis* | -2.51 | 586 |
| *Ardea alba* | 5.41 | 4994 |
| *Ardea cinerea* | 4.32 | 5303 |
| *Arenaria interpres* | 17.86 | 1696 |
| *Aythya ferina* | 11.34 | 744 |
| *Aythya fuligula* | 10.45 | 857 |
| *Aythya marila* | 0.43 | 577 |
| *Aythya nyroca* | 9.77 | 1444 |
| *Botaurus stellaris* | 17.73 | 182 |
| *Branta bernicla* | 2.86 | 853 |
| *Branta leucopsis* | 1.69 | 4057 |
| *Branta ruficollis* | 1.10 | 539 |
| *Bubulcus ibis* | 22.96 | 1297 |
| *Bucephala clangula* | -1.11 | 420 |
| *Calidris alba* | 18.86 | 254 |
| *Calidris alpina* | 11.83 | 467 |
| *Calidris canutus* | 19.08 | 550 |
| *Calidris maritima* | -2.48 | 498 |
| *Calidris minuta* | 22.77 | 697 |
| *Charadrius alexandrinus* | 19.20 | 630 |
| *Charadrius hiaticula* | 22.11 | 1103 |
| *Ciconia ciconia* | 22.95 | 707 |
| *Clangula hyemalis* | -2.50 | 2258 |
| *Cygnus columbianus* | 2.44 | 5300 |
| *Cygnus cygnus* | -1.49 | 1796 |
| *Cygnus olor* | 1.27 | 5837 |
| *Egretta garzetta* | 21.26 | 65 |
| *Fulica atra* | 5.86 | 2653 |
| *Fulica cristata* | 22.59 | 4535 |
| *Gallinula chloropus* | 5.61 | 862 |
| *Gallinago gallinago* | 18.58 | 944 |
| *Gavia arctica* | 0.76 | 581 |
| *Gavia stellata* | 3.05 | 949 |
| *Grus grus* | 14.72 | 379 |
| *Haematopus ostralegus* | 15.34 | 549 |
| *Himantopus himantopus* | 22.13 | 4140 |
| *Larus minutus* | 3.70 | 172 |
| *Larus argentatus* | 4.58 | 3413 |
| *Larus audouinii* | 11.45 | 2046 |
| *Larus canus* | 1.62 | 207 |
| *Larus fuscus* | 18.58 | 1707 |
| *Larus genei* | 11.90 | 607 |
| *Larus marinus* | -2.49 | 4695 |
| *Larus melanocephalus* | 9.31 | 478 |
| *Larus ridibundus* | 6.58 | 596 |
| *Limosa lapponica* | 19.23 | 4323 |
| *Limosa limosa* | 21.01 | 3760 |
| *Marmaronetta angustirostris* | 5.95 | 87 |
| *Melanitta fusca* | 0.67 | 904 |
| *Melanitta nigra* | 2.28 | 838 |
| *Mergellus albellus* | -1.58 | 2396 |
| *Mergus merganser* | -0.40 | 3945 |
| *Mergus serrator* | -1.08 | 1911 |
| *Phalacrocorax pygmeus* | 2.74 | 286 |
| *Netta rufina* | 5.35 | 1208 |
| *Numenius arquata* | 18.89 | 1738 |
| *Numenius phaeopus* | 22.06 | 228 |
| *Nycticorax nycticorax* | 23.47 | 218 |
| *Oxyura leucocephala* | 1.27 | 168 |
| *Pelecanus crispus* | 8.37 | 116 |
| *Pelecanus onocrotalus* | 22.01 | 71 |
| *Phalacrocorax aristotelis* | 2.30 | 499 |
| *Phalacrocorax carbo* | 3.70 | 6160 |
| *Philomachus pugnax* | 22.96 | 465 |
| *Phoenicopterus roseus* | 20.58 | 440 |
| *Platalea leucorodia* | 17.95 | 192 |
| *Plegadis falcinellus* | 23.14 | 1269 |
| *Pluvialis apricaria* | 5.36 | 692 |
| *Pluvialis squatarola* | 18.84 | 742 |
| *Podiceps auritus* | 3.39 | 4742 |
| *Podiceps cristatus* | 3.25 | 798 |
| *Podiceps grisegena* | 2.27 | 1256 |
| *Podiceps nigricollis* | 6.63 | 253 |
| *Porphyrio porphyrio* | 8.20 | 1810 |
| *Rallus aquaticus* | 4.52 | 563 |
| *Recurvirostra avosetta* | 22.04 | 1094 |
| *Somateria mollissima* | -7.53 | 3114 |
| *Tachybaptus ruficollis* | 3.54 | 4917 |
| *Tadorna ferruginea* | 8.88 | 500 |
| *Tadorna tadorna* | 4.30 | 2444 |
| *Sterna sandvicensis* | 12.38 | 422 |
| *Tringa erythropus* | 21.03 | 473 |
| *Tringa nebularia* | 23.16 | 748 |
| *Tringa ochropus* | 21.10 | 1475 |
| *Tringa totanus* | 15.48 | 1412 |
| *Vanellus vanellus* | 4.52 | 2933 |

## Appendix 4: Additional monitoring information and CTI correction

The International Waterbird Census (IWC) started for some species in the 1960s, but had comprehensive species coverage by the end of the 1980s. To be cautious, we started the study period in 1993. However, in some countries gulls and shags were not included directly in the IWC. The full waterbird species census was performed later in Romania (1999), Belgium (Flandre, 2000), Denmark (2001), United Kingdom (2002), Ireland (2002) and Sweden (still not full). As a change in species monitored can artificially affect the community changes, we took these dates into account in the analyses.

The community temperature index (CTI) was corrected to account for the monitoring changes in countries where the full waterbird species census started after the beginning of the study period (countries listed above). In these countries, the CTI values before the year(s) of monitoring change were centred per site (not reduced) and added to the average site CTI value of the years after the monitoring change. Hence, the addition of new species after the monitoring change doesn’t strongly affect the CTI values (Appendix 4, Table S1). Note that under the hypothesis of a CTI increase over years, the CTI correction leads to an overestimation of the site CTI average before the monitoring change. Regarding the CTI_sd_ no adaptation was done.

**Table S1:**
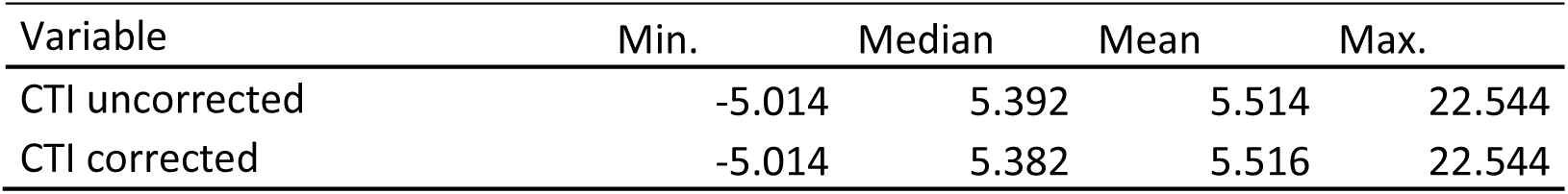
Summary of the variance minimum (Min.) median, mean and maximum (Max.) between the original CTI computed without and with correction.

| Variable | Min. | Median | Mean | Max. |
| --- | --- | --- | --- | --- |
| CTI uncorrected | -5.014 | 5.392 | 5.514 | 22.544 |
| CTI corrected | -5.014 | 5.382 | 5.516 | 22.544 |

We performed models with the full dataset and the data subset to control the potential differences. We used the same model framework as in the Methods section to evaluate the change of CTI, CTI_sd_, number of cold-dwelling species and number of warm-dwelling species. As a result, the models outputs were fairly similar between the two dataset, at the exception that warm-dwelling species did not significantly increased more than cold-dwelling species inside PAs (Appendix 4, Table S2).

**Table S2:** Comparison of the models with the full dataset and the subset of data with the full species monitoring. Protected area (PA) effect on temporal trends of the community temperature index (CTI) and standard deviation of the CTI (CTI_sd_), number of cold- and warm-dwelling species. Base line is sites outside PA and cold-dwelling species. Years were standardized to zero mean (std.) in the thermal-dwellers model and interactions are notified by ‘:’.

| Dataset | Variable | Parameter | Coefficient | SE | Z-value | P-value |
| --- | --- | --- | --- | --- | --- | --- |
| Full dataset | CTI | Intercept | -6.030 | 1.696 | -3.554 | < 0.001 |
|  |  | Year | 0.006 | 0.001 | 6.676 | < 0.001 |
|  |  | PA | 7.972 | 2.435 | 3.274 | < 0.001 |
|  |  | Year:PA | 0.004 | 0.001 | 3.412 | < 0.001 |
|  | CTI<sub>sd</sub> | Intercept | 4.294 | 1.602 | 2.680 | 0.007 |
|  |  | Year | 0.000 | 0.001 | 0.360 | 0.719 |
|  |  | PA | 11.310 | 2.300 | 4.920 | < 0.001 |
|  |  | Year:PA | 0.006 | 0.001 | 5.066 | < 0.001 |
|  | Cold- and Warm-dwellers | Intercept | 1.466 | 0.012 | 125.070 | < 0.001 |
|  |  | Year(std.) | 0.057 | 0.002 | 33.560 | < 0.001 |
|  |  | PA | 0.163 | 0.017 | 9.600 | < 0.001 |
|  |  | Dweller | 0.338 | 0.002 | 154.600 | < 0.001 |
|  |  | Year(std.):PA | 0.005 | 0.002 | 2.360 | 0.018 |
|  |  | Year(std.):Dweller | 0.008 | 0.002 | 3.820 | < 0.001 |
|  |  | PA:Dwellers | 0.022 | 0.003 | 7.360 | < 0.001 |
|  |  | Year(std.):PA:Dweller | 0.006 | 0.003 | 1.960 | 0.050 |
| Subset with full species monitoring | CTI | Intercept | 0.098 | 2.241 | 0.044 | 0.965 |
|  |  | Year | 0.003 | 0.001 | 2.563 | 0.010 |
|  |  | PA | 15.302 | 3.123 | 4.900 | < 0.001 |
|  |  | Year:PA | 0.008 | 0.002 | 4.930 | < 0.001 |
|  | CTI<sub>sd</sub> | Intercept | 2.396 | 2.051 | 1.168 | 0.243 |
|  |  | Year | 0.001 | 0.001 | 1.290 | 0.197 |
|  |  | PA | 8.743 | 2.858 | 3.059 | 0.002 |
|  |  | Year:PA | 0.004 | 0.001 | 3.153 | 0.002 |
|  | Cold- and Warm-dwellers | Intercept | 1.473 | 0.012 | 124.750 | < 0.001 |
|  |  | Year(std.) | 0.049 | 0.002 | 26.090 | < 0.001 |
|  |  | PA | 0.166 | 0.017 | 9.720 | < 0.001 |
|  |  | Dweller | 0.334 | 0.002 | 141.280 | < 0.001 |
|  |  | Year(std.):PA | 0.008 | 0.003 | 3.060 | 0.002 |
|  |  | Year(std.):Dweller | 0.014 | 0.002 | 6.020 | < 0.001 |
|  |  | PA:Dwellers | 0.024 | 0.003 | 7.460 | < 0.001 |
|  |  | Year(std.):PA:Dweller | 0.005 | 0.003 | 1.620 | 0.104 |

## Appendix 5: Protected area surfaces in the study area. Protected area surface (km²) is represented by points located at the centre of the corresponding cell (5°×5° resolution), which include both protected and not protected sites and at least 15 sites. The protected area surface corresponds to the sum of the PA surfaces per cell. The size of the points indicates the protected area surface size

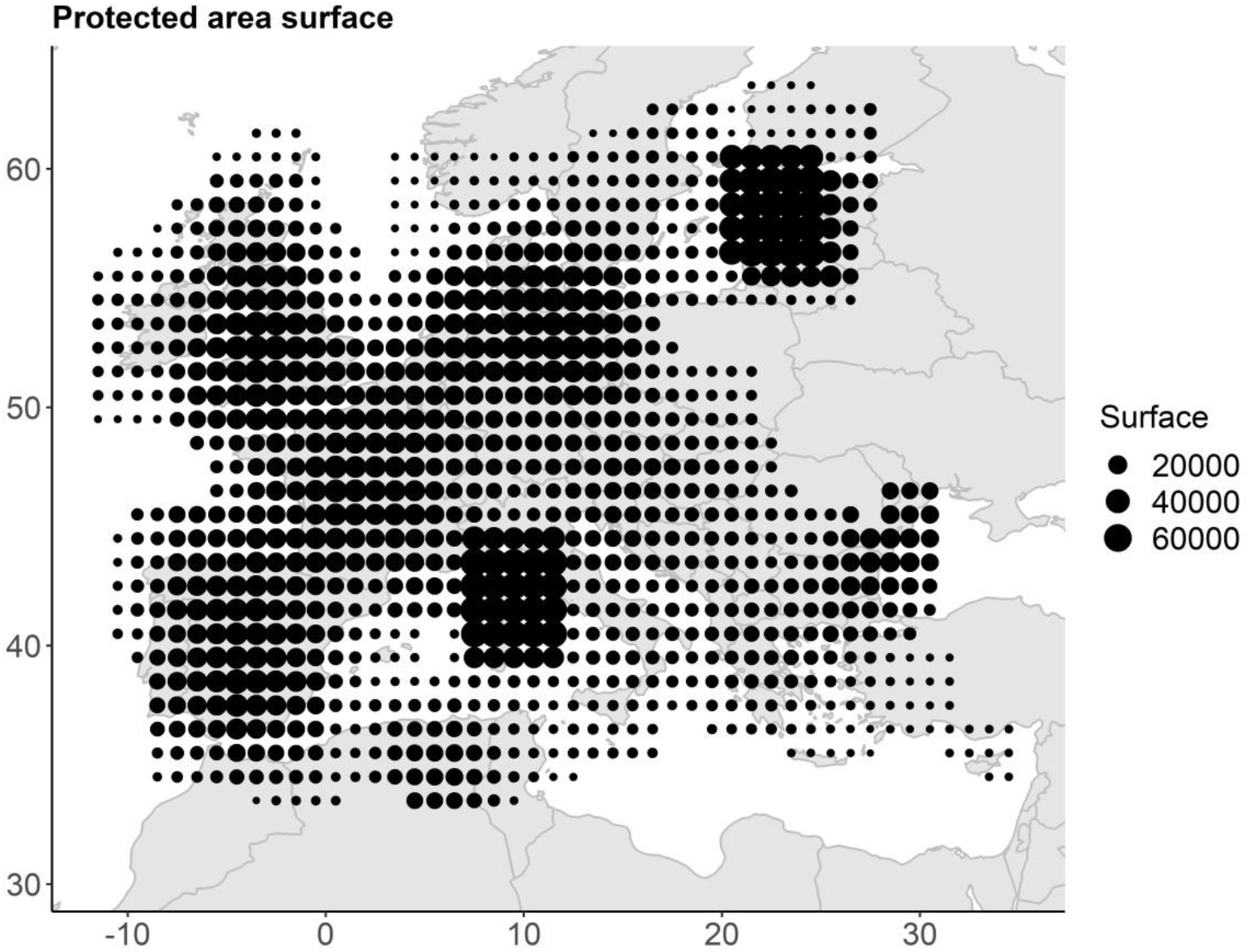

